# Structural insights into GlcNAc-1-phosphotransferase that directs lysosomal protein transport

**DOI:** 10.1101/2021.03.25.436948

**Authors:** Shuo Du, Guopeng Wang, Zhiying Zhang, Chengying Ma, Ning Gao, Junyu Xiao

**Author notes:** Correspondence: Ning Gao, Junyu Xiao. These authors contributed equally to this work (Shuo Du, Guopeng Wang).

## Abstract

GlcNAc-1-phosphotransferase (GNPT) catalyzes the initial step in the formation of the mannose-6-phosphate tag that labels ∼60 lysosomal proteins for transport. Mutations in GNPT cause lysosomal storage disorders such as mucolipidoses. However, the molecular mechanism of GNPT remains unclear. Mammalian GNPTs are α2β2γ2 hexamers in which the core catalytic α- and β-subunits are derived from GNPTAB. Here, we present the cryo-electron microscopy structure of the *Drosophila melanogaster* GNPTAB homolog (DmGNPTAB). Four conserved regions located far apart in the sequence fold into the catalytic domain, which exhibits structural similarity to that of the UDP-glucose glycoprotein glucosyltransferase (UGGT). Comparison with UGGT revealed a putative donor substrate-binding site, and the functional requirements of critical residues in human GNPTAB were validated using *GNPTAB*-knockout cells. DmGNPTAB forms an evolutionarily conserved homodimer, and perturbing the dimer interface undermines the maturation and activity of human GNPTAB. These results provide important insights into GNPT function and related diseases.

## Introduction

Protein phosphorylation is universally present as a regulatory strategy in eukaryotic cells. Glycan phosphorylation, though not as abundant, also plays essential roles in modulating cellular processes, particularly within the secretory compartments. For example, two novel secretory pathway glycan kinases have been recently characterized: the proteoglycan xylose kinase Fam20B regulates the biosynthesis of chondroitin sulfate and heparan sulfate proteoglycans (Koike et al., 2009; Wen et al., 2014; Zhang et al., 2018), and the protein O-mannose kinase POMK/SgK196 monitors the proper glycosylation of α-dystroglycan (Walimbe et al., 2020; Yoshida-Moriguchi et al., 2013; Zhu et al., 2016).

A canonical glycan phosphorylation event involves ∼60 secretory proteins that are destined for lysosomes. Similar to other secretory molecules, these lysosomal proteins are first synthesized in the endoplasmic reticulum and then traverse through the Golgi network. At the Golgi apparatus, these proteins are “phosphorylated” on a terminal mannose residue in their N-linked glycans, resulting in the formation of a mannose 6-phosphate (M6P) tag that is recognized by two specific M6P receptors to direct their lysosomal transport. Interestingly, this essential modification is not performed by an ATP-dependent kinase but is generated by the sequential action of two enzymes: first, the N-acetylglucosamine-1-phosphotransferase (GNPT) catalyzes the addition of an N-acetylglucosamine-1-phosphate (GlcNAc-1-P) group to the terminal mannose, and the GlcNAc-1-phosphodiester α-N-acetylglucosaminidase (NAGPA) then removes the GlcNAc moiety to uncover M6P (Varki and Kornfeld, 2015).

Mammalian GNPTs are large protein complexes that comprise two α-subunits, two β-subunits and two γ-subunits (Bao et al., 1996; Kudo and Canfield, 2006). The catalytic α- and β-subunits are first synthesized as GNPTAB fusion proteins. GNPTAB has a complex structural organization and includes two transmembrane (TM) segments, four conserved regions (CR1-CR4), two Notch repeats (N1 and N2), a DNA methyltransferase-associated protein (DMAP) interaction domain, and several spacer regions (S1-S4) (Figure 1A). The two TM segments anchor GNPTAB on the Golgi membrane and project most of the molecule into the Golgi lumen. The four CRs are conserved in Stealth proteins, and members of this family can function as hexose-phosphate transferases in bacteria to synthesize cell wall polysaccharides (Sperisen et al., 2005). The Notch repeats and DMAP interaction domain mediate the interaction between GNPT and its diverse lysosomal protein substrates (Qian et al., 2013; Qian et al., 2015; van Meel et al., 2016). The spacer regions are also functionally important. For example, the S2 region is responsible for interacting with the γ-subunit (De Pace et al., 2015; Velho et al., 2016). S3 contains a recognition site for the site-1 protease, which cleaves GNPTAB into its α- and β-subunits and thereby leads to catalytic activation (Marschner et al., 2011). The S1 spacer facilitates proper processing of GNPTAB by the site-1 protease (Liu et al., 2017). S4 contains an EF-hand calcium-binding motif, but its function remains unclear. The γ-subunit is encoded by *GNPTG* and plays a regulatory function to promote the activity of the GNPT holoenzyme toward a subset of substrates (Qian et al., 2010).

**Figure 1.**
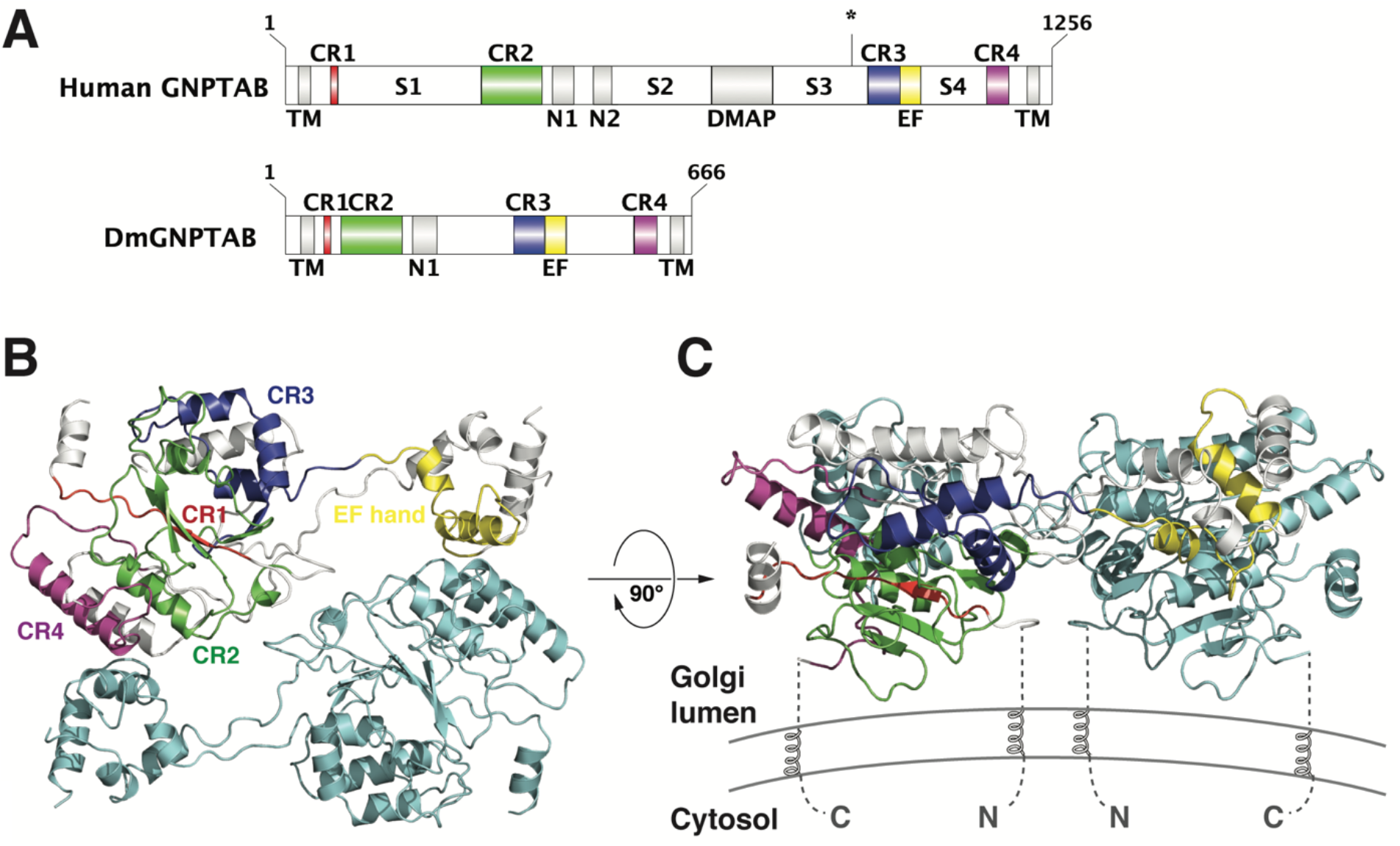
Cryo-EM structure of DmGNPTAB. **A**. Domain architectures of human GNPTAB and DmGNPTAB. TM: transmembrane segment; CR1-CR4: conserved regions in the Stealth proteins; N1 and N2: Notch repeats; DMAP: DNA methyltransferase-associated protein interaction domain; S1-S4: spacer regions. The asterisk indicates the S1P cleavage site in human GNPTAB. **B**. Ribbon diagram of the DmGNPTAB structural model. The four CRs and EF hand motif in one protomer are depicted in red, green, blue, magenta, and yellow, respectively; whereas the rest of this protomer is shown in white. The other protomer is shown in cyan. **C**. Putative model of DmGNPTAB on the Golgi membrane. The N- and C-termini of each molecule are indicated.

More than 200 mutations in GNPTAB have been documented in patients with mucolipidoses, a group of human lysosomal storage disorders that include skeletal and neuronal abnormalities (summarized in Velho et al., 2019). Mutations in GNPTAB and GNPTG have also been linked to stuttering (Frigerio-Domingues and Drayna, 2017). On the other hand, GNPT might serve as a potential antiviral target because a number of viruses, such as the Ebola virus and the common cold coronaviruses OC43 and 229E, rely on the lysosomal pathway for infection and egress (Flint et al., 2019; Wang et al., 2021). However, the underlying molecular mechanism of GNPT remains insufficiently understood. Here, we sought to characterize the structural architecture of GNPT by cryo-electron microscopy (cryo-EM) and successfully determined the 3.5-Å structure of the *Drosophila melanogaster* GNPTAB. Our results reveal critical structural features that are conserved in the GNPTAB family. We then generated a *GNPTAB*-knockout cell line using the CRISPR-Cas9 genome editing technique and validated the importance of residues in human GNPTAB involved in donor substrate binding and dimerization. We also analyzed pathogenic missense mutations and assessed their potential impacts. Together, our results advance the understanding of GNPT and related human diseases.

## Results

### Cryo-EM structure of *Drosophila melanogaster* GNPT

We sought to investigate the structural basis of GNPT function. Despite intensive attempts, we were unable to obtain the structure of the GNPT complex and could not determine the structure of the α/β subcomplex or the GNPTAB precursor. *Drosophila melanogaster* has a GNPTAB homolog (DmGNPTAB) but lacks a discernable gene encoding GNPTG (Sperisen et al., 2005; Tiede et al., 2005). The DmGNPTAB protein is markedly more compact than its human counterpart (Figure 1A). Specifically, this protein contains all four CRs and a similar CR3-CR4 spacer but has shorter CR1-CR2 and CR2-CR3 spacers, only one Notch repeat, and lacks the DMAP interaction domain and S1P cleavage site (Figure 1A, Figure S1). We obtained the luminal portion of DmGNPTAB and analyzed its structure by cryo-EM (Figure 1B, Figure S2). The structure was determined at an overall resolution of 3.5 Å (Table S1). The center region displayed high resolutions, which allowed us to build the structural model de novo. Approximately half of the protein molecule, including all four CRs and the CR3-CR4 spacer, could be confidently placed. The amino and carboxyl termini are located on the same side of a monomer, which sheds light on the topology of the full-length protein on the Golgi membrane (Figure 1C). The rest of the molecule, particularly the CR2-CR3 spacer including the Notch repeat, displayed weak densities and was thus not modeled.

### Structure of the DmGNPTAB monomer

Two domains are clearly visible in the DmGNPTAB monomer (Figure 2A). The large domain, which is referred to as the Stealth domain, features an α/β fold that includes all four Stealth CRs. CR1 contains a β-strand that occupies the center of the Stealth domain. CR2 comprises three β-strands and two α-helices. The three strands sandwich the CR1 strand in a parallel manner to form a β-sheet, whereas the two helices are situated on each side of the sheet. CR3 consists of one strand and three helices. The single CR3 strand runs antiparallel to the four strands described above, and the three helices bundle together with the short helix in CR2. CR4 features a single helix, which packs against the long helix in CR2. The small domain is encoded by the CR3-CR4 spacer, including the calcium-binding EF hand motif, and features a helix bundle to mediate dimerization.

**Figure 2.**
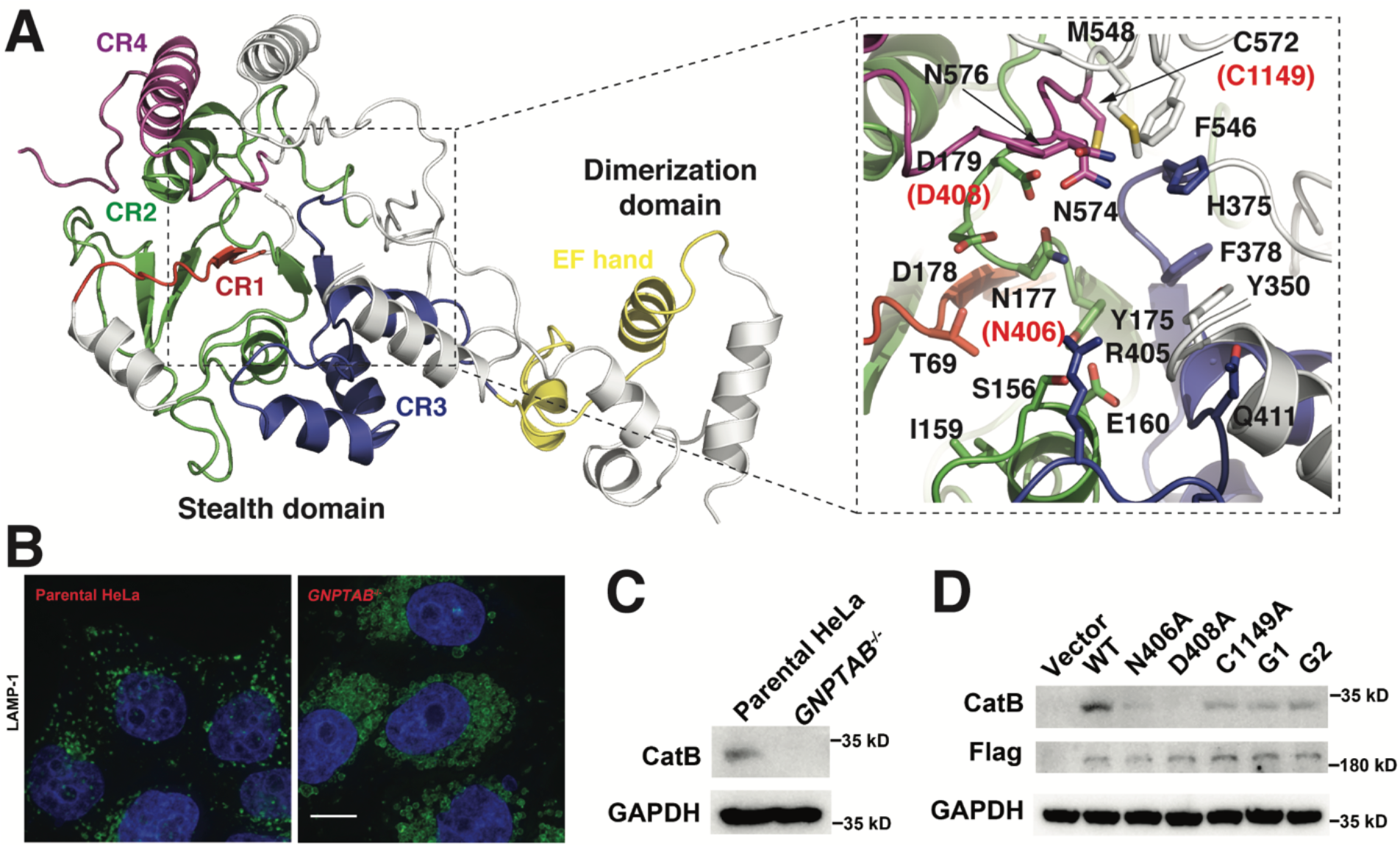
Putative UDP-GlcNAc-binding pocket. **A**. DmGNPTAB contains a large Stealth domain and a small dimerization domain. Residues involved in forming the putative UDP-GlcNAc-binding pocket are highlighted in the enlarged panel. Human GNPTAB residues N406, D408, and C1149 that correspond to N177, D179, and C572 in DmGNPTAB are highlighted red. **B**. Confocal immunofluorescence images of parental and GNPTAB-deficient HeLa cells stained using the late endosomal/lysosomal marker LAMP-1 (green). Nuclei are stained with DAPI (blue). The scale bar is 10 μm. **C**. Mature cathepsin B (CatB) was not detected in GNPTAB-deficient cells. **D**. GNPTAB mutants could not fully rescue CatB processing in GNPTAB-deficient cells. The expression of GNPTAB was monitored using anti-Flag antibody.

The four Stealth CRs, which are located far apart in the sequence, fold into a single globular domain. The Stealth domain shows structural similarities to several GT-A-type glycosyltransferases (Lairson et al., 2008) despite low sequence homology (Figure S3A). In particular, this domain exhibits structural resemblance to UDP-glucose glycoprotein glucosyltransferase (UGGT), particularly at the central β-sheet region (Figure S3B). UGGT is an ER-resident protein that surveils the folding status of secretory glycoproteins (Braakman and Bulleid, 2011). Similar to GNPT, UGGT is a label marker, and the labeling of UGGT also occurs on the terminal mannose of an N-linked glycan. Instead of labeling proteins that are destined for lysosomes, UGGT labels proteins that are incompletely folded: specifically, it recognizes misfolded glycoproteins and transfers a glucose residue to the terminal mannose on the glycan of these proteins, and this modification is recognized by ER chaperones, including calnexin and calreticulin, to facilitate correct folding.

Crystal structures of UGGT homologs from several thermophilic fungi have been determined (Roversi et al., 2017; Satoh et al., 2017). A structural comparison between the Stealth domain of DmGNPTAB and the catalytic domain of *Thermomyces dupontii* UGGT (TdUGGT) in complex with UDP-glucose offers a glimpse into the sugar nucleotide-binding site of GNPT. UDP-glucose binds to the surface pocket of TdUGGT (Figure S3B). DmGNPTAB has a similar surface pocket, which is formed by a group of conserved residues, including Thr69 from CR1; Ser156, Ile159, Glu160, Tyr175, Asn177, Asp178, and Asp179 from CR2; His375, Phe378, Arg405, and Gln411 from CR3; Phe546 and Met548 from the CR3-CR4 spacer; and Cys572, Asn574, and Asn576 from CR4 (Figure 2A). The functional importance of these residues is underscored by the fact that missense mutations of a number of the corresponding human residues have been found in patients (see Discussion). It is likely that the donor substrate UDP-GlcNAc is accommodated in DmGNPTAB in an orientation similar to that of UDP-glucose in TdUGGT, with the left side of the pocket enclosing the uridine diphosphate moiety and the right side holding the N-acetylglucosamine group (Figure S3B). A Ca^2+^ ion facilitates the accommodation of UDP-glucose in TdUGGT and is coordinated by three Asp residues, including Asp1294 and Asp1296 in the Asp-X-Asp signature motif of GT-A glycosyltransferases and Asp1427 (Figure S3C). A well-conserved Asn177-Asp178-Asp179 motif in CR2 of DmGNPTAB aligns with the Asp-X-Asp motif in TdUGGT, whereas Cys572 appears to take the position of TdUGGT-Asp1427. It is thus likely that these residues are also involved in the binding to a divalent cation that assists in positioning the UDP-GlcNAc in DmGNPTAB.

### Human GNPTAB mutants are functionally defective

We sought to validate the functional importance of some of these residues in human GNPTAB. To unambiguously analyze the activities of various GNPTAB mutants, we first generated a *GNPTAB*-knockout (*GNPTAB*^*-/-*^) HeLa cell line using the CRISPR/Cas9 genome-editing technique (Figure S4). Consistent with previous observations (van Meel et al., 2016), *GNPTAB*^*-/-*^ cells exhibited markedly swollen lysosomes compared with the parental cells (Figure 2B), which indicated that lysosomal function is severely impaired in these cells. As a result of lysosomal dysfunction, the lysosomal cysteine protease cathepsin B (CatB) was not properly processed, and the mature form of endogenous CatB at ∼30 kDa was not detected in the lysates of *GNPTAB*^*-/-*^ cells (Figure 3C). This finding was also consistent with previous observations obtained with GNPTAB-deficient HAP1 cells (Flint et al., 2019). We then generated alanine substitutions of Asn406, Asp408, and Cys1149, which are equivalent to Asn177, Asp179, and Cys572 in DmGNPTAB that form the putative metal-binding site (Figure S1, Figure S3C), and examined the abilities of these mutants to rescue CatB maturation. As expected, the expression of wild-type (WT) GNPTAB restored the mature form of CatB (Figure 2D). In contrast, N406A, D408A, and C1149A displayed reduced activities compared with the WT protein, and D408A appeared to be completely inactive. These results demonstrate the importance of these residues for GNPT function, corroborating our structural analyses.

**Figure 3.**
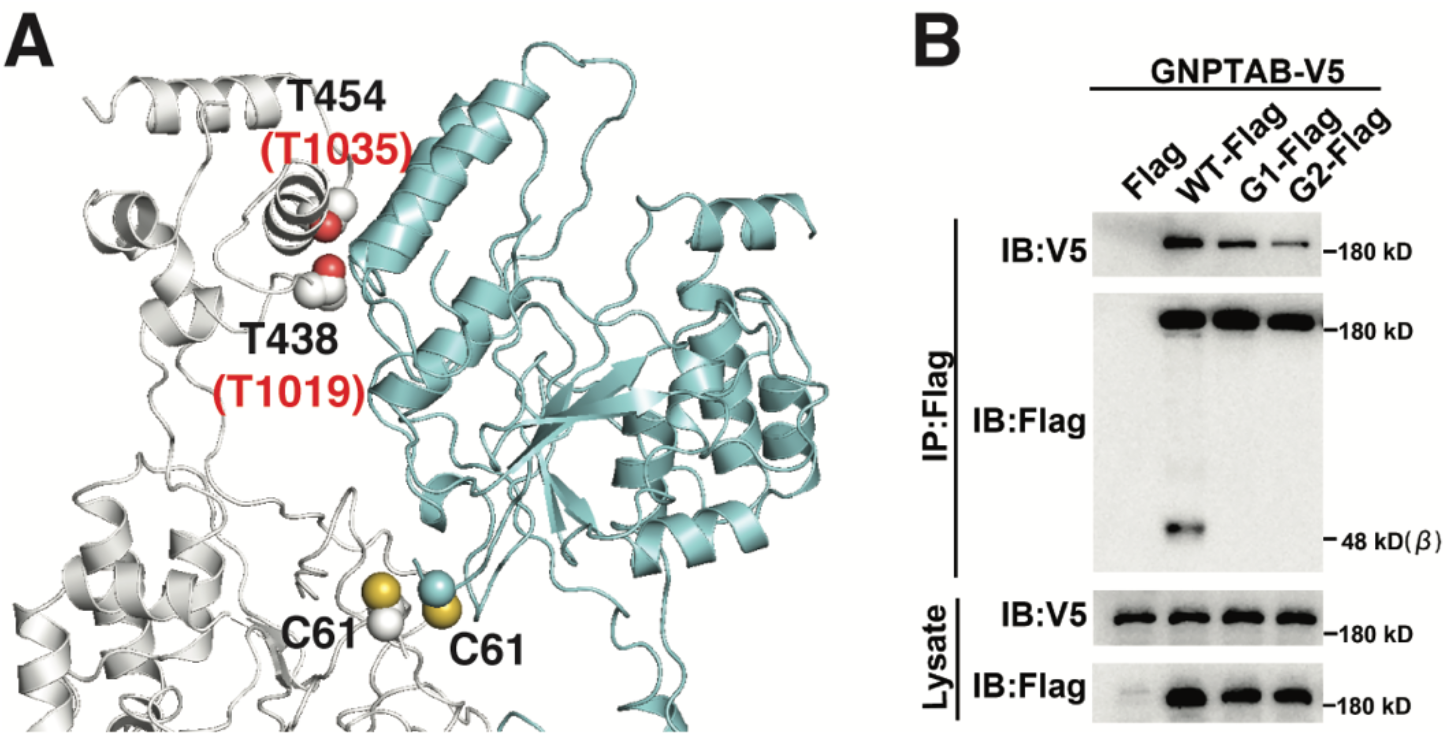
Mutations in the dimer interface perturb human GNPTAB dimerization and maturation. **A**. Structure of the DmGNPTAB dimer. Cys61, Thr438, and Thr454 are highlighted. **B**. G1 (T1019N/D1020G/Q1021S) and G2 (T1035N/R1036G/I1037S) displayed defective processing and diminished abilities to form heterodimers with WT GNPTAB.

### A conserved dimeric architecture

Mammalian GNPTs are α2β2γ2 hexameric complexes, and the construction of the hexamer remains poorly understood. In our structure, two DmGNPTAB molecules form a homodimer that resembles two fishes nestling against each other in a head-to-tail orientation (Figure 1B). The dimer is mainly mediated by the CR3-CR4 spacer and CR4, whose counterparts both reside in the β-subunit of human GNPT (Figure 1A). A few residues in CR2 also contribute to dimer formation. The dimer interface buries ∼1,500-Å^2^ solvent-accessible surfaces from each molecule and involves a number of invariant residues (Figure S1), which suggests that the dimerization mechanism observed in this study is likely generally conserved in the GNPTAB family.

To verify the functional relevance of the dimer, we generated two human GNPTAB mutants: G1 (T1019N/D1020G/Q1021S) and G2 (T1035N/R1036G/I1037S). These two mutants were designed to create sites that allow the attachment of N-linked glycans. Thr1019 and Thr1035 in human GNPTAB correspond to Thr438 and Thr454 in DmGNPTAB, both of which are located in the dimer interface (Figure 3A). The introduction of bulky glycans at these positions would impede dimerization of the human protein. We tagged these mutants with Flag tags at the C-terminus, co-expressed them with V5-tagged WT GNPTAB in HEK293T cells, and performed Flag immunoprecipitation. Flag-tagged and V5-tagged WT GNPTAB proteins were efficiently coprecipitated (Figure 3B), which indicated the formation of GNPTAB dimers in the cells. In contrast, the interactions between G1 or G2 and WT GNPTAB were markedly reduced, which suggested that these two mutants exhibit decreased abilities to dimerize with the WT protein. Furthermore, unlike WT GNPTAB, these two mutants were not efficiently processed because the ∼48-kDa band that corresponds to the β-subunit was not observed after their expression (Figure 3B). Importantly, these mutations could also not fully rescue the maturation of CatB in *GNPTAB*^*-/-*^ cells (Figure 2D). Together, these results demonstrated that proper dimer formation is important for the processing of GNPTAB and for the activity of the GNPT holoenzyme.

## Discussion

More than 200 mutations in GNPTAB, including at least 65 missense mutations, have been detected in patients with various forms of mucolipidoses (Velho et al., 2019). The effects of some of these mutations have been characterized biochemically, but the structure of human GNPTAB has remained elusive despite its long research history. The structure of DmGNPTAB allows extrapolation of the core structure of human GNPT and can thus be used to further assess the molecular impacts of disease-causing mutations (Figure 4). For example, the human GNPTAB residues Ser385, Glu389, Asp407, Asp408, His956, Arg986, and Asn1153 correspond to Ser156, Glu160, Asp178, Asp179, His375, Arg405, and Asn576 in DmGNPTAB (Figure 4A), all of which are found among the sugar nucleotide-binding pockets described above (Figure 2A). Thus, their mutations likely affected the binding of UDP-GlcNAc, consistent with the findings from previous studies showing that missense mutations involving these sites, such as S385L, E389K, D407A, D408N, H956Y, R986C, and N1153, resulted in markedly decreased enzyme activities (Danyukova et al., 2020; Ludwig et al., 2017; Qian et al., 2015). We also showed that D408A was unable to restore the maturation of CatB in *GNPTAB*^*-/-*^ cells (Figure 2D). Another set of mutants, including W81L, R334Q and R334L, S399F, I403T, and D1018G, displayed defects in exit from the ER, which indicated protein misfolding (Qian et al., 2015). These sites were also conserved in DmGNPTAB (Trp70, Arg105, Ser170, Leu174, and Asp437; Figure 4A). Trp70 and Leu174 are both present in the interior of the structure and are intimately packed with surrounding hydrophobic residues. Arg105 forms a bidentate interaction with Glu607 to support Pro601, which is involved in dimerization. Ser170 likely forms hydrogen bond interactions with Asp65 and Arg115, and the corresponding human residues, Asp76 and Arg344, are also mutated in some patients (Figure 4A). Similarly, some of the other mutations also lead to structural disturbance of human GNPT, as can be rationalized by the DmGNPTAB structure.

**Figure 4.**
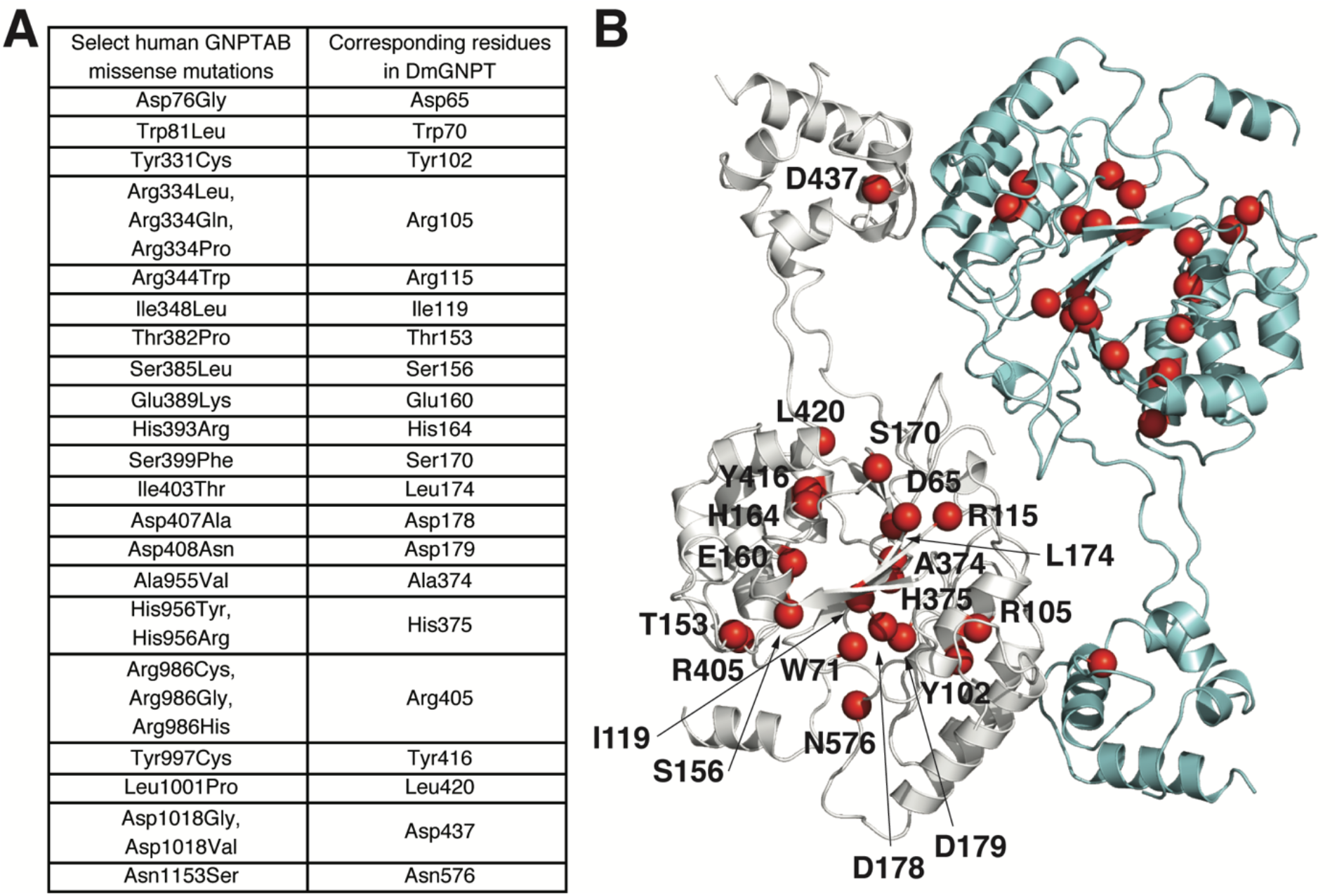
Insights into human mucolipidoses. **A**. Select human GNPTAB missense mutations and the corresponding residues in DmGNPTAB. **B**. The positions of DmGNPTAB residues listed in the previous panel are shown in the structure.

Human GNPT is an α2β2γ2 hexamer. The DmGNPTAB structure reveals a dimeric framework that sheds light on the assembly mechanism of human GNPT. The CR3-CR4 spacer and CR4 play dominant roles in mediating the formation of the DmGNPTAB homodimer, which likely reflects how the β2 dimer is formed in GNPT because most of the residues involved in the dimer interface are conserved (Figure S1) and because mutations of two of these residues reduced human GNPTAB dimerization (Figure 3B). In addition to the interactions between β-subunits, Cys70 in human GNPTAB is involved in disulfide-linked homodimerization of the α-subunits (De Pace et al., 2015). Cys61 in DmGNPTAB appears to align with Cys70, and in the DmGNPTAB dimer, two Cys61 residues from the two monomers are located in close proximity (Figure 3A), which suggests that a disulfide bond between the corresponding Cys70s could be readily formed to further stabilize the α2β2 subcomplex. The γ-subunit also forms a disulfide-linked homodimer (Kudo and Canfield, 2006), and only a γ2 dimer can be assembled into the GNPT holoenzyme (Encarnacao et al., 2011). The γ2 dimer, through its interactions with the S2 spacers in the two α-subunits, supplies another layer of interaction to architect the α2β2γ2 hexamer.

The most critical issue regarding the molecular mechanism of GNPT, namely, its specific targeting of 60 lysosomal proteins among thousands of glycoproteins that traverse the Golgi apparatus, remains to be addressed. We were unable to model the sole Notch repeat domain in DmGNPTAB due to its weak density, which indicates its structural flexibility in the absence of a cognate acceptor substrate. The α-subunit of human GNPT contains two Notch repeats and a DMAP interaction domain, all of which are involved in binding to lysosomal proteins, although their relative contribution differs among each individual substrate (Qian et al., 2015; van Meel et al., 2016). Furthermore, the γ-subunit, which is absent in *Drosophila* and contains an M6P receptor homology domain and another DMAP interaction domain, also plays an important role in the activity of GNPT toward select lysosomal proteins (Qian et al., 2010). It is apparent that starting from an ancestral Stealth gene, GNPTAB family proteins have gained increasing complexity in eukaryotes and even obtained additional interacting partners in vertebrates to ensure the proper transport of lysosomal proteins.

In summary, we have elucidated the structure of DmGNPT, and our findings offer a more in-depth understanding of the function of human GNPTAB and provide molecular insights into the pathogenesis of human mucolipidoses caused by GNPTAB mutations.

## Materials and Methods

### Protein expression and purification

The *Drosophila melanogaster GNPTAB* gene (CG8027) was cloned from a fly cDNA library. The DNA fragment encoding DmGNPTAB residues 50-630 was cloned into the psMBP2 vector (Tagliabracci et al., 2016), which facilitates its expression in insect cells as a secreted His6-MBP fusion protein with a tobacco etch virus (TEV) protease cleavage site. Bacmids were generated in DH10Bac cells using the Bac-to-Bac system (Invitrogen). Sf21 insect cells grown in SIM SF medium (Sino Biological Inc.) were used to generate and amplify the baculoviruses. For protein production, Hi5 cells grown in SIM HF medium (Sino Biological Inc.) were infected at a density of 1.5–2.0 × 10^6^ cells/ml. Forty-eight hours later, conditioned media were collected by centrifugation at 2,000 g for 30 min. The media were then concentrated using a Hydrosart Ultrafilter (Sartorius) and transferred into binding buffer containing 25 mM Tris-HCl, pH 8.0, and 150 mM NaCl. The recombinant protein was then isolated using Ni-NTA resin (GE healthcare) and eluted with a buffer containing 25 mM Tris-HCl, pH 8.0, 150 mM NaCl, and 500 mM imidazole. After TEV protease digestion for 10 h at 4°C to remove the N-terminal His6-MBP tag, the protein mixture was transferred into a buffer containing 25 mM Tris-HCl, pH 8.0, and 50 mM NaCl using a Centricon with a cutoff of 10 kDa (Millipore). The untagged proteins were then purified by anion exchange chromatography (Resource Q) and eluted using a 50–1000 mM NaCl salt gradient in 25 mM Tris-HCl, pH 8.0, followed by size exclusion chromatography (Superdex Increase 200) and elution in 25 mM HEPES, pH 7.5, and 150 mM NaCl.

### Cryo-EM data collection, model building and structure analyses

Four-microliter aliquots of purified DmGNPTAB at 0.5 mg/ml were applied onto glow-discharged Quantifoil holey-carbon grids (R1.2/1.3, 300 mesh, gold), blotted at 4 °C in 100% humidity, and plunged into liquid ethane with a Vitrobot Mark IV (FEI). The cryogrids were screened with a 200-kV Talos Arctica microscope. Data collection was performed with a 300-kV Titan Krios G3 microscope equipped with a Gatan GIF Quantum K2 Summit direct electron detector using SerialEM (Mastronarde, 2005). The statistics for data collection and processing are summarized in Table S1.

Movie frames were aligned with MotionCor2 (Zheng et al., 2017). The CTF parameters were estimated using Gctf (v1.063) (Zhang, 2016). Micrographs were sorted based on image qualities, and 14,345 micrographs were used for subsequent reconstruction with Relion (v3.07) (Zivanov et al., 2018). Particles were autopicked based on the templates generated by manual picking and subjected to 2D classification. Classes showing clear structural details were then used to create an initial model. 3D classifications with C1 symmetry were first performed, and 544,196 particles with good structural features were selected for further 3D classifications using C2 symmetry. A total of 131,301 particles were then selected for 3D refinement, which resulted in a map with an overall resolution of 3.53 Å after Bayesian polishing and CTF refinement. The resolution was estimated using the gold-standard Fourier shell correlation (FSC) 0.143 criteria.

The structure of DmGNPTAB was built de novo in Coot (Emsley et al., 2010). Bulky aromatic residues and N-linked glycosylation sites were used as landmarks during the structural modeling process. Structure refinement was performed using real-space refinement in Phenix (v1.18) (Adams et al., 2010). A structural homology search was performed using the DALI server (Holm, 2020). Figures were prepared with ESPript (Gouet et al., 2003), PyMOL (Schrödinger), and UCSF Chimera (Pettersen et al., 2004).

### Generation and characterization of *GNPTAB*^***-/-***^ **HeLa cells**

HeLa cells were grown in Dulbecco’s modified Eagle’s medium (DMEM) supplemented with 10% (v/v) fetal bovine serum (FBS) at 37°C in a 5% CO2 incubator. *GNPTAB*^*-/-*^ cells were generated using CRISPR/Cas9 technology. The guide RNA, 5’-ACAAAACATGGTATTGATCT-3’, which targets exon 2 of *GNPTAB*, was cloned into the pSpCas9(BB)-2A-GFP vector (Addgene, 48138). Three micrograms of the plasmid was then transfected into HeLa cells using Lipofectamine 2000 (Thermo Scientific). Two days after transfection, GFP-positive cells were sorted into single clones using an Astrios EQ cell sorter (Beckman Coulter). Single clones were cultured in 96-well plates for two weeks. The genome type of the knockout cells was determined by DNA sequencing.

For the visualization of lysosomes, parental and GNPTAB^-/-^ HeLa cells were fixed with 4% paraformaldehyde in phosphate-buffered saline (PBS) at room temperature for 15 min, permeabilized with PBS containing 0.1% saponin and 2% bovine serum albumin, and then incubated with anti-LAMP1 antibody (Santa Cruz, sc-20011, 1:50) at 4°C overnight. The next day, the cells were washed with PBS and incubated with Alexa Fluor 488 donkey anti-mouse antibody (Invitrogen, A21202, 1:150) for 1 h at 25°C. The coverslips containing the cells were then re-washed with PBS, incubated with DAPI for the visualization of nuclei, and examined with a Delta Vision microscope. The images were analyzed using Velocity (v6.1.1) software.

To examine the maturation of CatB, WT and mutant GNPTAB-Flag plasmids were transfected into *GNPTAB*^*-/-*^ HeLa cells as indicated. Seventy-two hours later, the cells were harvested, washed with PBS, lysed in lysis buffer (PBS supplemented with 0.5% Triton X-100, 1% protease inhibitor cocktail, and 1 mM phenylmethylsulfonyl fluoride) for 15 min at 4°C, and analyzed by western blotting.

### Immunoprecipitation and western blotting

The human *GNPTAB* gene was cloned into modified pcDNA 3.1 vectors that encode a C-terminal Flag or V5 tag. Mutations were generated using a PCR-based method and verified by sequencing. For the immunoprecipitation experiments, V5-tagged GNPTAB was cotransfected with Flag-tagged GNPTAB into HEK293T cells in 10-cm dishes using polyethyleneimine. Forty-eight hours later, the cells were harvested, washed with PBS, and lysed in lysis buffer. The lysates were then centrifuged for 15 min at 13,000 g, and the supernatant was incubated with Flag M2 beads (Sigma, A2220) for 2 h at 4°C. The beads were washed three times with lysis buffer. The immunoprecipitated proteins were eluted from the beads using the 3X Flag peptide (NJPeptide, NJP50002) and analyzed by western blotting.

For the western blot analyses, protein samples separated by SDS-PAGE were transferred to nitrocellulose membranes, and the membranes were then blocked with 4% nonfat milk for 30 min at 25°C and incubated with primary antibody overnight at 4°C. The next day, the membranes were incubated with HRP-conjugated secondary antibodies in 4% nonfat milk for 1 h at room temperature. Detections were performed by enhanced chemiluminescence using an Amersham Imager 800. The primary antibodies used for immunoblotting were anti-Flag (ABclonal, AE005, 1:1000), anti-Flag (MBL, PM020, 1:1000), anti-V5 (Santa Cruz, sc-81594, 1:1000), anti-V5 (Millipore, AB3792, 1:1000), anti-CatB (Cell Signaling, 31718, 1:1000), and anti-GAPDH (TransGen, HC301-02, 1:5000). The secondary antibodies were goat anti-mouse (TransGen, HS201-01, 1:5000) and goat anti-rabbit (TransGen, HS101-01, 1:5000).

## Acknowledgments

We thank the Core Facilities at the School of Life Sciences, Peking University for help with negative-staining EM; the Cryo-EM Platform of Peking University for help with data collection; the High-performance Computing Platform of Peking University for help with computation; the National Center for Protein Sciences at Peking University for assistance with cell sorting. The work was supported by the National Key Research and Development Program of China (2020YFC0848700 and 2017YFA0505200 to J.X., 2019YFA0508904 to N.G.), the National Science Foundation of China (31822014 to J.X., 31725007 and 31630087 to N.G.), and the Qidong-SLS Innovation Fund to J.X. and N.G..

## Author contributions

S.D. performed most of the experiments. S.D. and G.W. prepared the cryo-EM sample, collected data, and processed cryo-EM images under the supervision of N.G. and J.X. J.X. built the structural model, analyzed the data, and wrote the manuscript with inputs from all authors.

## Data and materials availability

The cryo-EM map and atomic coordinates of DmGNPTAB have been deposited in the EMDB and PDB with accession codes EMD-30910 and 7DXI, respectively.

**Figure S1.**
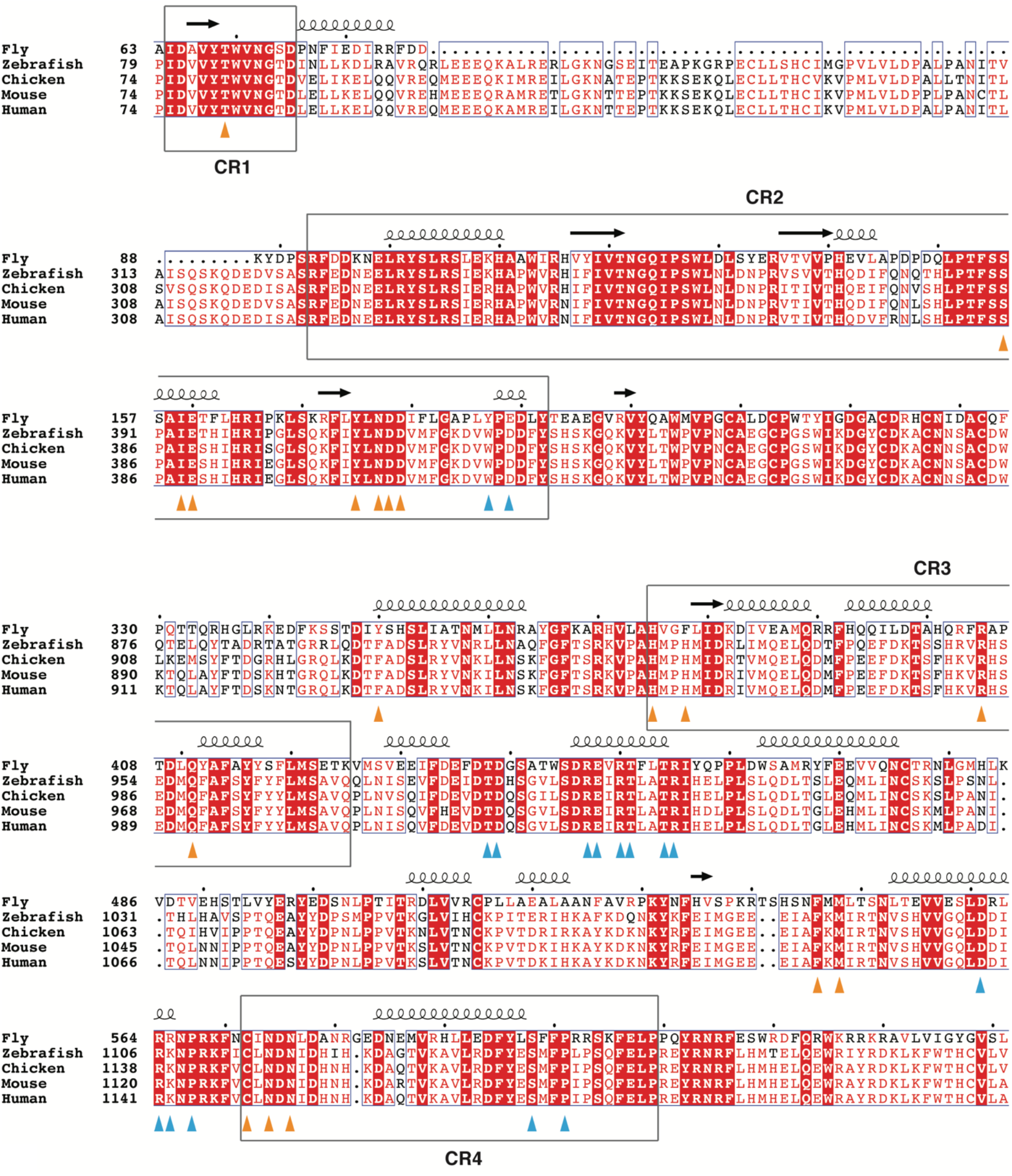
Structure-based sequence alignment of select GNPTAB homologs around the four CRs. The residues that are likely involved in sugar nucleotide binding and dimer formation are indicated by orange and cyan triangles below the sequence blocks, respectively.

**Figure S2.**
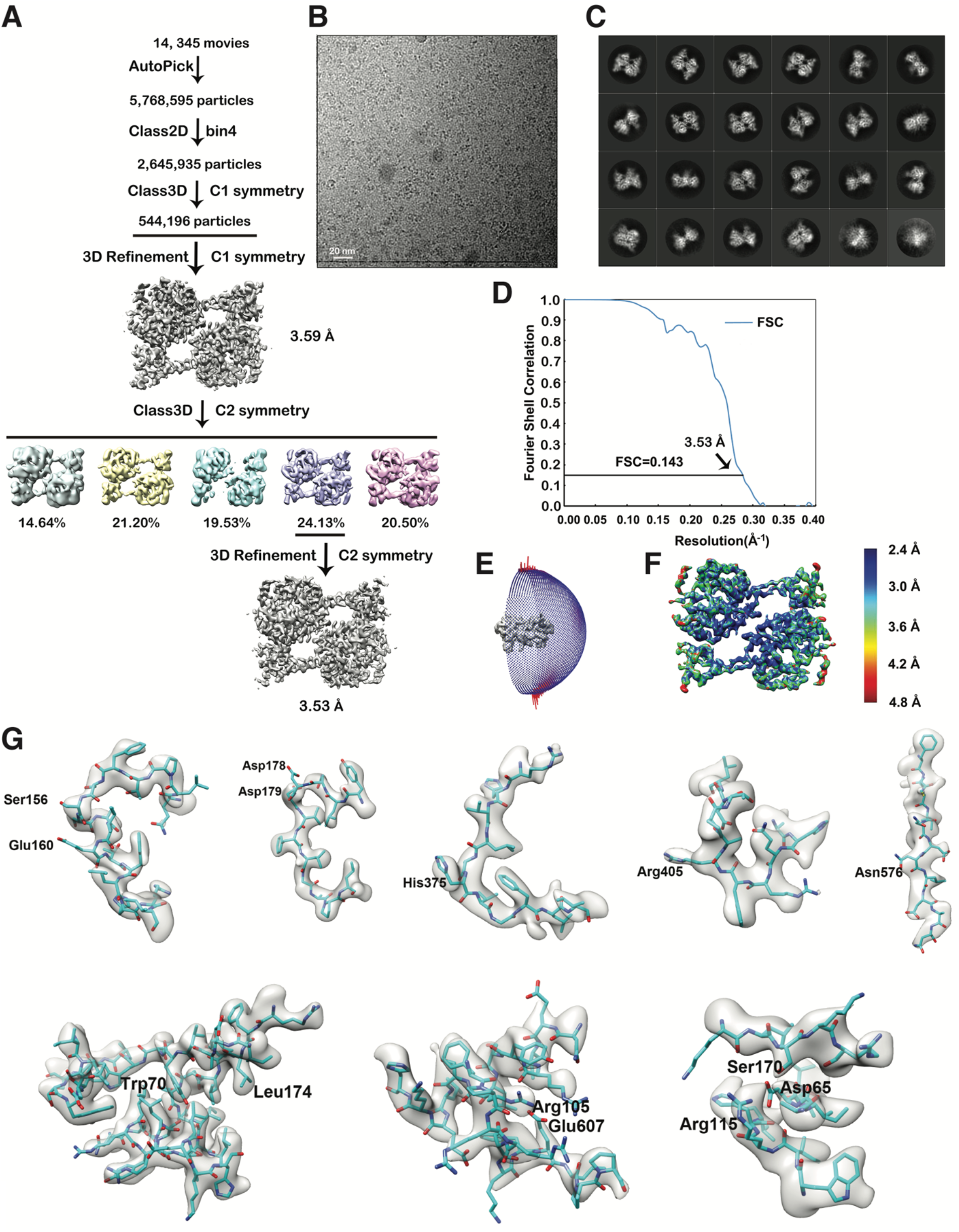
Workflow for cryo-EM structure reconstruction. **A**. Workflow for cryo-EM data processing. **B**. Representative raw cryo-EM image. **C**. Representative 2D classes. **D**. Gold-standard Fourier shell correlation (FSC) curve with estimated resolution. **E**. Eulerian angle distribution of the particles used in the final 3D refinement. **F**. Local resolution estimation of the final map. **G**. Cryo-EM density maps around select DmGNPTAB residues described in the manuscript.

**Figure S3.**
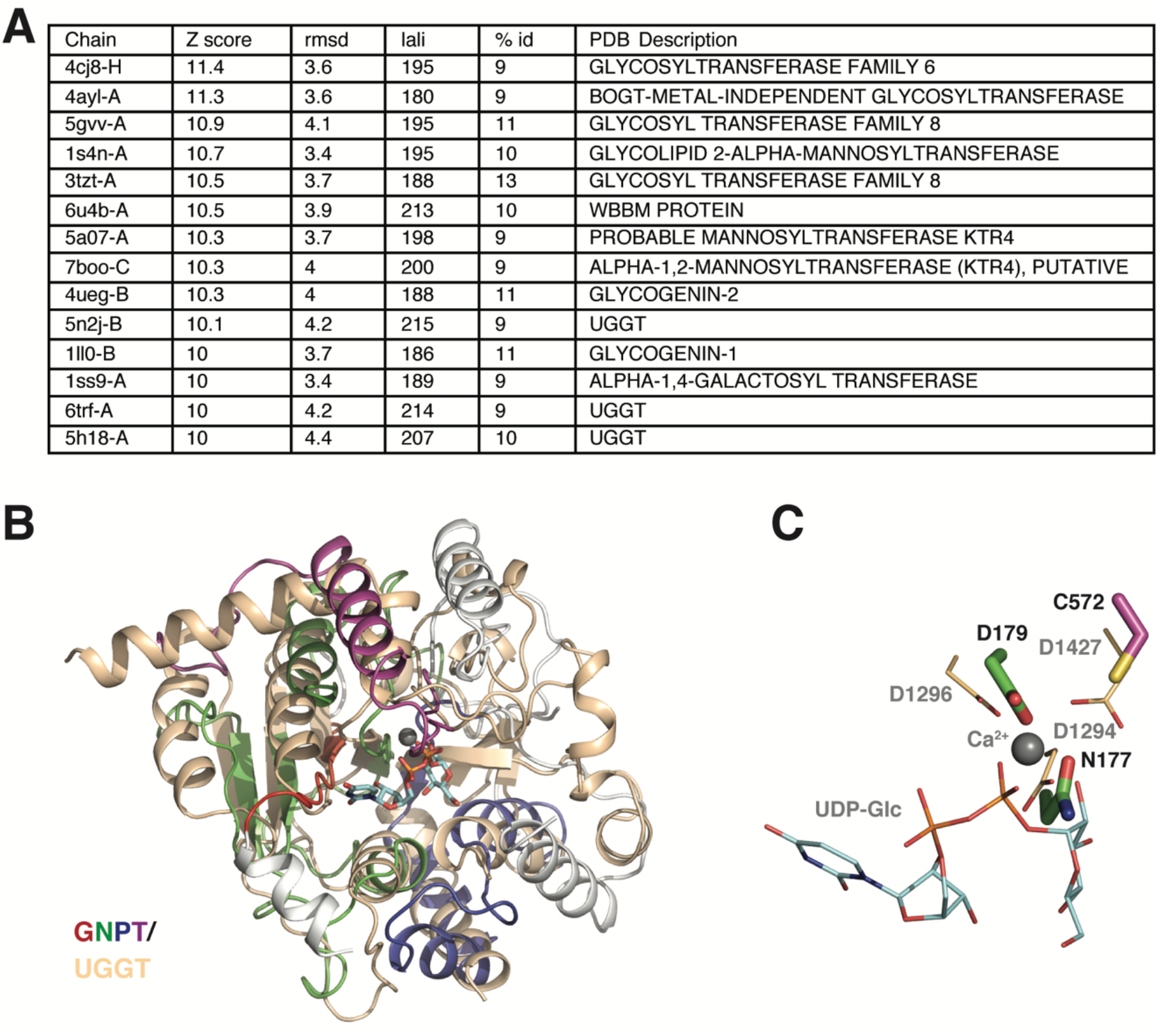
The DmGNPTAB Stealth domain displays structural homology to a number of glycosyltransferases, including UGGT. **A**. Top hits from a DALI search against a nonredundant subset of PDB structures. **B**. Structural superposition of DmGNPTAB and TdUGGT (PDB ID: 5H18). Both structures are shown using the same color scheme as in Figure 2. **C**. Asn177, Asp179, and Cys572 in DmGNPTAB appear to align with Asp1294, Asp1296, and Asp1427 in TdUGGT, which coordinate a Ca^2+^ ion to position UDP-Glc.

**Figure S4.**
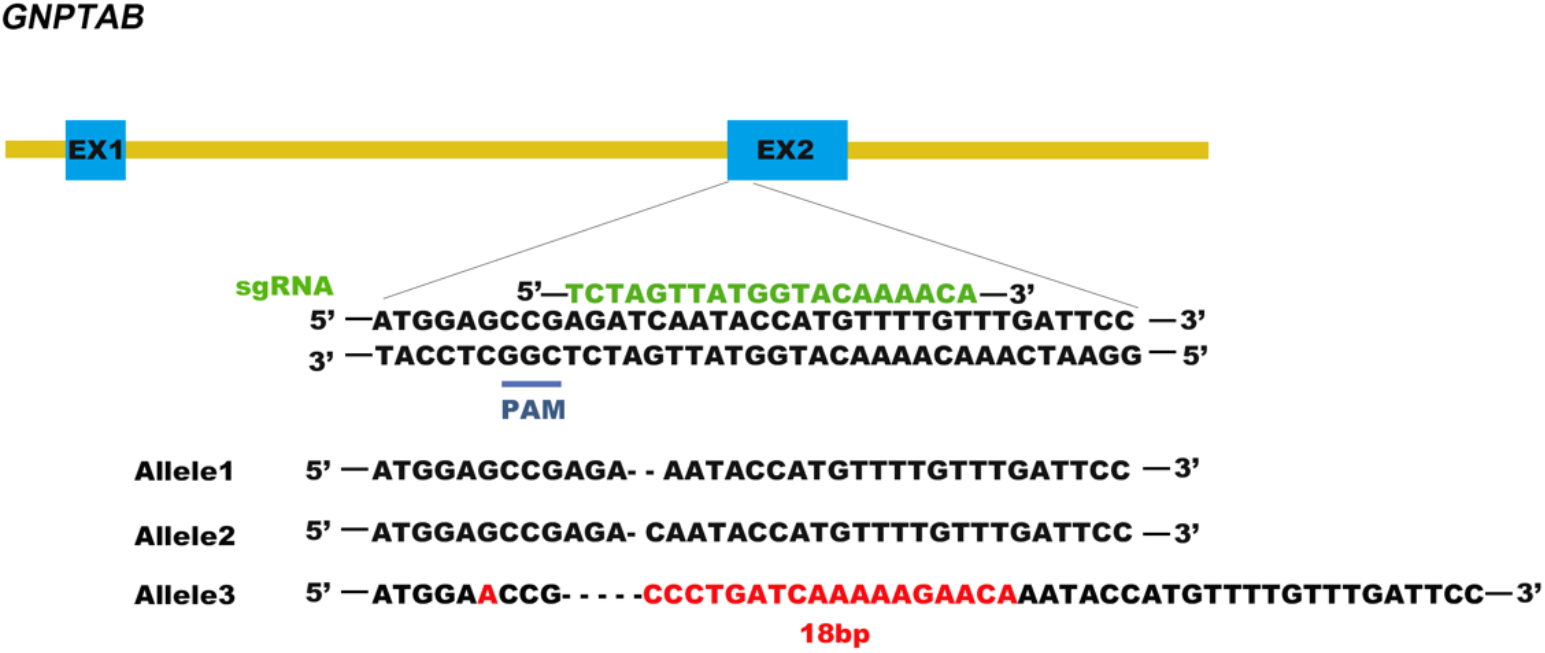
Generation of the *GNPTAB*-knockout HeLa cell line by CRISPR-Cas9 genome editing. The sequences of the mutated alleles are shown.

**Table S1.** Cryo-EM data collection, processing and validation statistics.

|  | GNPTAB (EMDB & PDB IDs: ) |
| --- | --- |
| **Data collection and processing** |  |
| Voltage (kV) | 300 |
| Microscope | FEI Titan Krios G3 |
| Camera | K2 Summit (Gatan) |
| Magnification (calibrated) | 210,000X |
| Electron exposure (e<sup>-</sup>/Å<sup>2</sup>) | 60.29 |
| Exposure rate (e<sup>-</sup>/Å<sup>2</sup>/s) | 18.84 |
| Number of frames collected per micrograph | 32 |
| Energy filter slit width | 20 eV |
| Automation software | SerialEM |
| Defocus range (μm) | -1.0 to -2.2 |
| Pixel size (Å) | 0.6516 |
| Micrographs used | 14,345 |
| Estimated accuracy of rotations | 2.447 |
| Symmetry imposed | C2 |
| Initial particle images | 5,768,795 |
| Final particle images | 131,301 |
| Resolution at 0.143 FSC of masked reconstruction (Å) | 3.53 |
| **Refinement** |  |
| Initial model used (PDB code) | none |
| Refinement package | Phenix v1.18.2 (Real-space refinement at 3.53 Å) |
| Map-model CC |  |
| CC_mask | 0.81 |
| CC_box | 0.73 |
| CC_peaks | 0.64 |
| CC_volume | 0.79 |
| Model composition |  |
| Non-hydrogen atoms | 6,130 |
| Protein residues | 742 |
| Ligands | 2 |
| *B* factors (Å<sup>2</sup>) |  |
| Protein | 24.82 |
| Ligands | 62.76 |
| R.m.s. deviations |  |
| Bond lengths (Å) | 0.006 |
| Bond angles (°) | 1.292 |
| Validation |  |
| MolProbity score | 1.89 |
| Clashscore | 4.71 |
| Poor rotamers (%) | 0.30 |
| Ramachandran plot |  |
| Favored (%) | 85.54 |
| Allowed (%) | 13.91 |
| Disallowed (%) | 0.55 |
| Cβ outliers (%) | 0.00 |

